# RACLET: the Ramp Above Critical Level Endurance Test to evaluate critical force in isometric task

**DOI:** 10.1101/2025.07.11.664330

**Authors:** Batiste Morel, Maximilien Bowen, Mylène Vonderscher, Leo Blervaque, Hervé Di Domenico, Julien Pernot, Pierre Samozino

**Affiliations:** Laboratoire Interuniversitaire de Biologie de la Motricité LIBM, EA 7424, Université Savoie Mont-Blanc, Le Bourget-du-Lac, France; Clinical research department, Centre Hospitalier Métropole Savoie, Chambéry, France; Centre Hospitalier Métropole Savoie, Centre de Réadaptation Respiratoire, 49 avenue du Grand Port 73100 Aix-les-Bains, France

**Author notes:** This research has been funded by the Agence Nationale de la Recherche [ANR-22-CE14-0073].

**Keywords:** Critical intensity, Muscle contraction, Fatigability All-out, Time-to-exhaustion

## Abstract

The Ramp Above Critical Level Endurance Test (RACLET) is a novel submaximal test designed to evaluate the parameters of the critical force model without strenuous exercise. This study aimed to validate the RACLET in healthy and pathological populations and to assess its reliability, concurrent validity, and predictive capacity. Sixteen healthy volunteers and ten patients with respiratory pathologies participated in this study. The RACLET consisted of a decreasing ramp force starting at 60% and ending at 15% of the maximum force for a total duration of 425 s with brief regular maximal voluntary contractions. The goodness of fit for the RACLET model on the maximal contraction force was excellent in both populations (median r^2^ *≈* 0.95). In patients, RACLET parameters demonstrated excellent reliability (ICC *>* 0.90). The concurrent validity of the critical force estimate compared with the all-out method was high (error: −0.3*±*7.4%). The model’s predictive capacity for time-to-exhaustion and fatigue during constant-intensity exercise was excellent (r^2^ = 0.910 and 0.907, respectively). The RACLET provides a reliable and valid estimate of critical force model parameters opening up numerous practical applications in vulnerable populations, individualised physical activity programs, and longitudinal monitoring. The feasibility and performance of the test make it a promising tool for assessing muscle function in various contexts.

## Introduction

Isometric force evaluation is the most widely used method for characterising muscle function. This is an experimentally convenient method with inexpensive sensors. Muscular strength is thus assessed in contexts as varied as athlete strength and conditioning training (Warneke et al., 2023), injury aftercare (Bisciotti et al., 2017), characterisation of chronically ill patients (Robles et al., 2011), and as a readout for innovative therapies (El Mhandi and Bethoux, 2013). Most often, maximum muscle strength is measured in the fresh state. However, this represents only one aspect of muscular function that is decoupled from endurance capacity, i.e. maintaining force over time. Therefore, the duration of sustained submaximal force is also regularly used to investigate muscular endurance, for example, in pathologies such as Chronic Obstructive Pulmonary Disease (COPD; Evans et al., 2015; Serres et al., 1998).

However, this approach has one major limitation: the endurance time depends on the level of force demanded, which is often determined (be it in absolute or relative values) without a strong rationale. More importantly, Vonderscher et al. (2024) showed that depending on the level of force chosen, opposite conclusions can be drawn. To overcome this crucial limitation, some studies used the critical intensity model (Chartogne et al., 2024; Leclercq et al., 2024; Veni et al., 2019).

The concept of critical intensity proposes a model that describes the relationship between exercise intensity and duration for which it can be sustained (Poole et al., 2016). The general form of this decreasing and converging relationship was identified during the second half of the 19th century (Haughton, 1873). This relationship was mathematically modelled almost a century later to characterise the muscular endurance capacity during isometric contractions (Monod and Scherrer, 1965; Rohmert, 1960). The duration *t*_*lim*_ that a force can be maintained before exhaustion can be expressed as

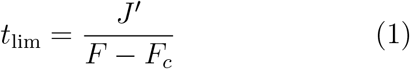

where *F* is the force produced (i.e. exercise intensity) and two constants: (i) the critical force *F*_*c*_ (in N), a threshold delimiting the efforts for which a physiologically stable state can be achieved from those where homeostasis cannot be preserved and a dramatic onset of fatigue is observed (Burnley et al., 2012; Burnley, 2022); (ii) a reserve of impulse, *J*′ (in N · s), which can be figuratively consumed above *F*_*c*_. Once emptied, it is no longer possible to maintain the required force *F* and a state of exhaustion is reached.

One of the major advantages of this approach is that it determines a hallmark of an individual’s exercise physiology independent of the test conditions. Moreover, *F*_*c*_ and *J*′ are related to distinct physiological and biomechanical mechanisms (Poole et al., 2016), enabling a better understanding of the determinants of muscular performance and accurate exercise prescription for effective interventions in health and training contexts (Meyler et al., 2025; Muniz and Meyler, 2025; Poole and Jones, 2023). However, a major obstacle to the widespread use of these indices is the difficulty of the required experimental procedures. The two most common methods are multiple time-to-exhaustion and all-out tests. The former generally consists of performing three to five efforts (> 24H intervals) at several constant force levels until exhaustion is reached. The data were then used to fit Eq. 1 and estimate the parameters *F*_*c*_ and *J*′. The latter consists of performing a single test of maximum sustained effort (usually 3–5 minutes). The converging force at the end of the test corresponds conceptually to the critical force *F*_*c*_, and the impulse above the end-test force corresponds to *J*′. For both methods, these strenuous sustained efforts are very difficult to implement in fragile populations, for the routine testing of athletes, or in a research context, when several experimental conditions must be tested.

Critical force represents a threshold in muscular physiology. It has sometimes been characterised as a “fatigability threshold” because fatigue (a reduction in maximum force capacity) progresses extremely rapidly above it but remains very limited below it (Burnley et al., 2012; Burnley, 2022). In fact, when the muscle is fatigued, it can recover when the intensity is below the critical force. Taking advantage of this characteristic, it would be possible to design a submaximal test (i.e. without reaching exhaustion) to identify the intensity at which the muscle switches from a state of fatigue to a state of recovery. A simple idea would be to start the test at an intensity above the critical force, which decreases over time and ultimately crosses the critical intensity (without knowing when *a priori*) which would allow for recovery. By regularly measuring the maximum force capacity during the test, it is possible to identify where the *fatigue*–*recovery* state transition occurs (i.e. the switch from a decrease to an increase in maximal capacity), corresponding to the critical force. Such a test can overcome the limits of feasibility of existing methods (*vide supra*).

Based on the pioneering work of H. Morton on the critical intensity framework (Hugh Morton, 1996; Morton and Billat, 2013), we recently proposed a mathematical model of exercise fatigability (Bowen et al., 2024). Interestingly, by observing fatigability (i.e. alteration in maximal capacity) during exercise, it was possible to determine the parameters of the critical intensity model. This offers a framework within which the proposed test can be developed and mathematically described. Thus, the aim of this study was to build a test called the Ramp Above Critical Level Endurance Test (RACLET), based on the model previously proposed by our team. This test is hypothesised to enable the evaluation of the parameters of the critical force model by inducing only moderate fatigue. Once theoretically developed, a knee extensor RACLET protocol was set up to test (i) the RACLET feasibility, (ii) the RACLET intersession reliability, (iii) the RACLET concurrent agreement with the all-out test, and (iv) the ability of RACLET to predict exercise fatigability and duration before exhaustion during constant-intensity exercise.

## Methods

### Model development

The change in maximal force capacity *F*_max_(*t*) during an exercise performed above the critical force *F*_*c*_ can be described by a linear function of the integral of the force above the critical force (hereafter referred as *J*_AC_) with 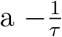 slope (Bowen et al., 2024). *Nota bene J*_AC_ converge toward a maximal value of impulse that can be realised above *F*_*c*_ being 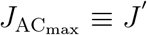. In addition, without fatigue, the initial force capacity is *F*_*i*_; therefore, the maximal force capacity at any time *F*_max_(*t*) is defined as

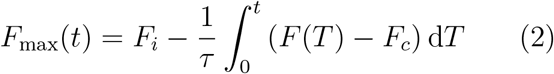

The RACLET is designed such that the intensity starts from a submaximal but supra-critical force (*F*_*c*_ *< F* ^*⋆*^ *≤ F*_*i*_) and decreases linearly with slope *S*:

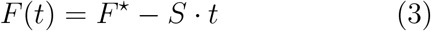

By combining Eq. 2 and Eq. 3, the maximal force capacity *F*_max_(*t*) evolves as a quadratic function of time,

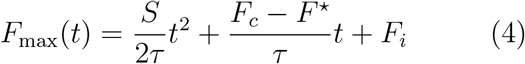

By developing a generic model (Eq. 2) for a decreasing ramp exercise, the theoretical decrease in the maximum force capacity *F*_max_(*t*) is described by a second-order polynomial function, which depends on three parameters: initial force *F*_*i*_ [N], critical force *F*_*c*_ [N], and typical time constant *τ* [s]. Eq. 4 can be written as *F*_max_(*t*) = *A · t*^2^ + *B · t* + *C*, where the three parameters (*A, B*, and *C*) are:

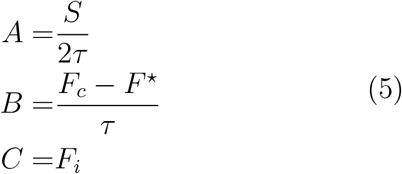

The theoretical evolution of the variables and a graphical representation of the parameters are shown in Fig. 1.

**Figure 1.**
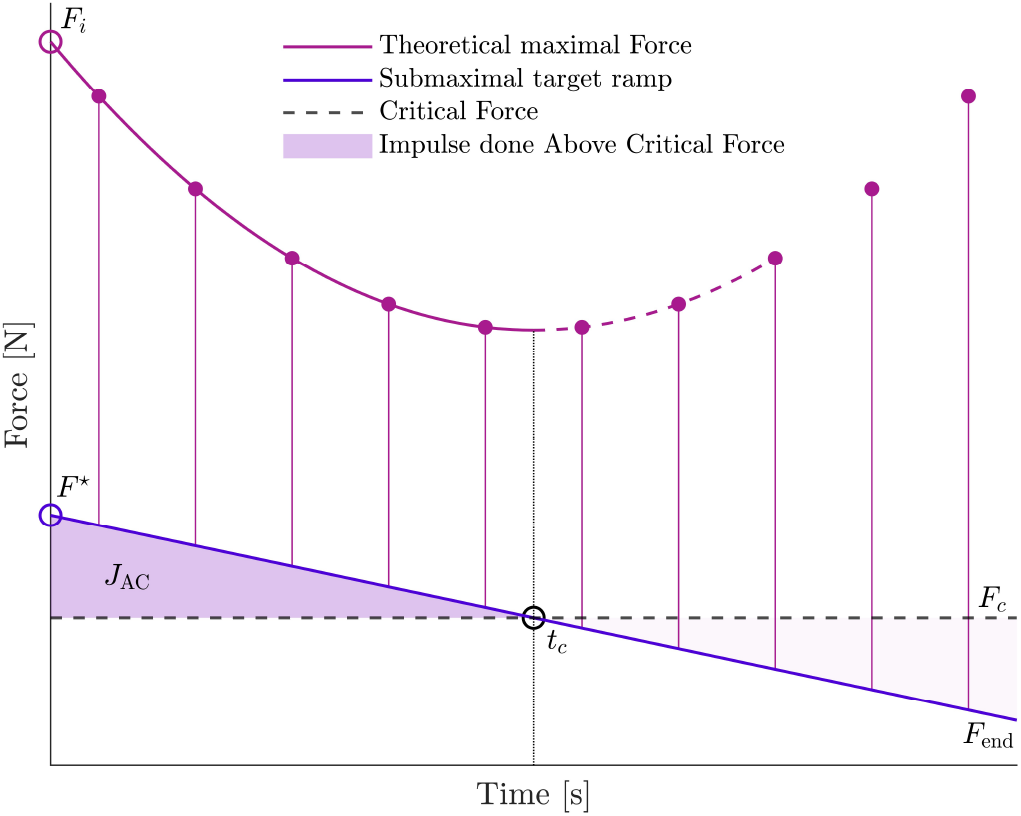
Theoretical representation of the Ramp Above Critical Level Endurance Test (RACLET). The blue line represents the target force starting at *F*\* and ending at *F*_*end*_. Regularly, the maximal force capacity is measured with maximal voluntary contraction (purple dot). During the first phase of the test, an impulse above the critical force *J*_*AC*_ is produced (shaded area). At this point, the participant is in a fatigue phase, and the maximal force, starting at *F*_*i*_, will decrease according to a theoretically quadratic law (purple line; Eq. 4). At an a priori unknown time *t*_*c*_, the target force will cross the critical force *F*_*c*_, and the participant will then start the recovery phase; the maximal force capacity will increase accordingly. For this phase, the model is displayed as a dotted purple line because it is not supposed to describe the recovery phase correctly.

### Participants and experimental design

This study involved two experiments. The first aimed to test the concurrent validity of the RACLET in a healthy and physically active population, and the second tested the feasibility and reliability of the RACLET in a population with respiratory pathology enrolled in a functional rehabilitation programme. Approval for this project was obtained from the local Ethics Committee on Human Experimentation. Written informed consent was obtained from all participants and the study was conducted in accordance with the Declaration of Helsinki.

For the first experiment, 16 volunteers participated in this study (5 females and 11 males; mean *±* s.d.; age: 22.8 *±* 3.2 years; mass: 70.5*±* 10.2 kg; height: 1.76 *±* 0.07 m). The experiment was conducted over five randomised sessions (one test per day) 24–48 h apart using intermittent isometric knee extension tasks. The experimental tests consisted of a RACLET, a 5-min all-out test, and three time-to-exhaustion tests. All participants were healthy and physically active, without any injuries, and free from any consumption of drugs, medications, or dietary supplements that could have influenced testing.

For the second experiment, 10 patients participated in this study (4 females and 6 males; pathology: 6 COPD, 3 chronic asthma, and 1 idiopathic fibrosis; mean *±* s.d.; age: 69.9 *±* 14.3 years; mass: 70.8 *±* 18.7 kg; height: 1.65 *±* 0.09 m). RACLET was performed by the clinical team as part of the patient’s functional evaluation at the beginning and end of the rehabilitation program (7 weeks apart). The patients did not present with any injuries in the lower limbs and realised the protocol within their routine treatments (including oxygen supplementation).

### Material (*Apparatus*)

The participants were seated upright in a custom-built chair with the right knee flexed at 90° and the hips at 100° (180° full extension; Fig. 2). The leg was attached 3 cm above the malleoli with a non-compliant strap to a strain gauge with its amplifier (Futek FSH04207; 444.822 N; M6×1; 500Hz Gain 1.9N/V). The force signals were sampled (200 Hz) and stored in a custom LabVIEW program (National Instruments Corporation, Austin, TX, USA) using a 16-bit analogue-to-digital interface Ni DAQ card (NI-USB6210 National Instruments Corporation).

**Figure 2.**
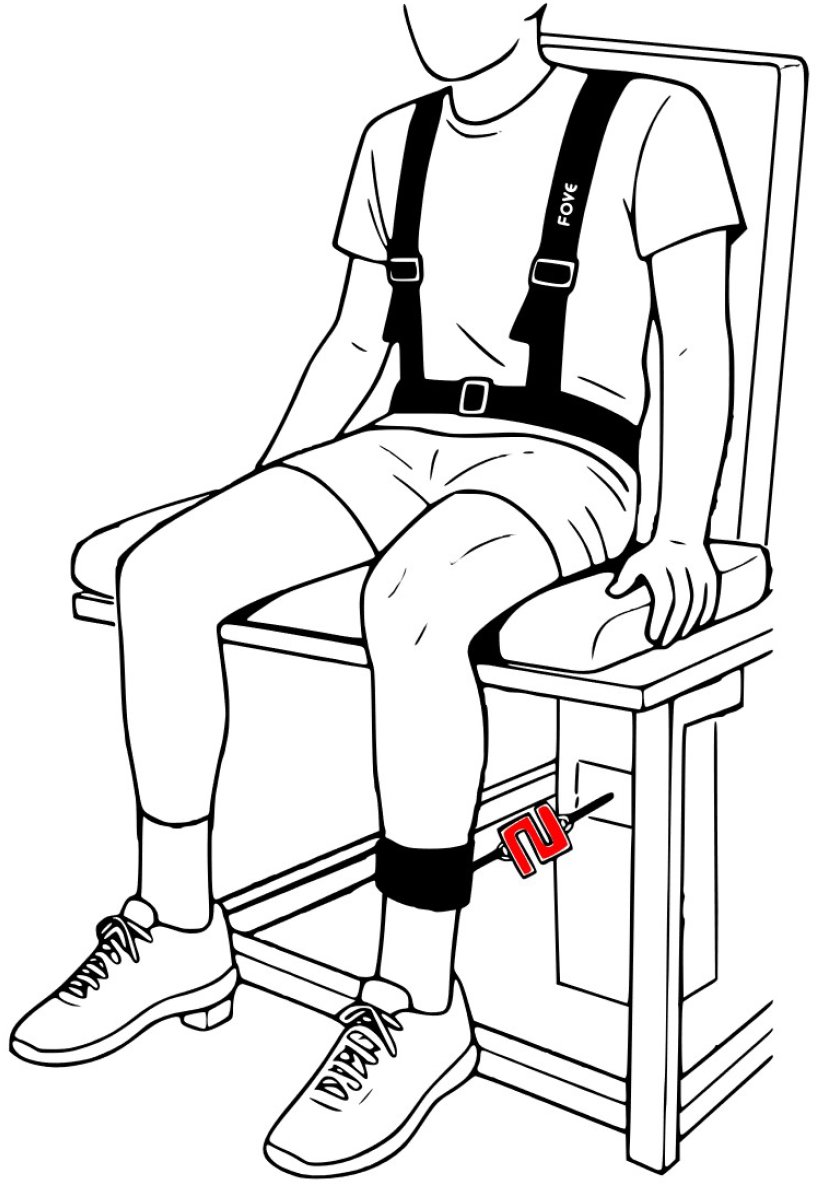
Custom-built chair for isometric knee extensors evaluation. Strain gauge, represented in red, was attached 3 cm above the malleoli so to measure the force applied by the knee extensors perpendicular to the leg.

### Methodology

To ensure respect for the force targets and duty cycle (3s on, 2s off for all tests), visual feedback was provided on the screen in direct vision and associated with audio signals. All maximum intensity efforts were supported by strong verbal encouragement from investigators or clinicians. Each experimental session began with a standardised 10-min warm-up (10, 8, 6, 4, and 2 contractions at 10, 20, 40, 60, and 80% of maximal perceived force, respectively, with 1-min recovery between sets). Then, two maximal voluntary contractions (MVC) were realised with a 3-min interval to evaluate the maximal isometric force (another MVC was performed if the difference was > 5%, the highest was recorded).

#### All-out test

The all-out test consisted of 60 MVCs and lasted for 300 s, with the maximum force of each contraction being recorded. To limit eventual pacing strategies, the participants were kept uninformed of the number of contractions completed and the remaining time left in the procedure. Furthermore, if the participant did not reach 95% of the maximal force recorded at the beginning of the experimental session (i.e. over the first three contractions), the test was stopped and a 3-min recovery period was observed before restarting the all-out test.

#### RACLET

The RACLET consisted of a decreasing ramp force starting at 60% and ending at 15% of the maximum force for a total duration of 425 s (85 contractions). Regularly (contraction numbers 5, 11, 18, 26, 35, 45, 55, 65, 75, and 85), the participants were asked to perform MVCs. Because the critical force of the knee extensors has been reported to be *≈* 25-35% MVC (Burnley, 2009), it was expected that the test would start above the critical force and end below it for all participants. In addition, the force ramp should reach the critical force two-thirds of the way through the test. In the first part of the test, the force of the MVCs should decrease (fatigue) and then increase (recovery) as the target force falls below the critical force. The rate of perceived exertion (RPE 6-20 scale) was measured immediately after each MVC, and the highest and end-test values were conserved.

#### Time-to-exhaustion tests

Each participant participated in three time-to-exhaustion tests. They had to maintain a randomly selected level of force between 45 and 65% of the maximal voluntary force, which was supposed to generate a duration of effort before exhaustion of 2 to 15 min. In addition, every 45 s (nine contractions), MVC was performed to analyse the development of fatigue prediction during the test. The test was stopped when the participant could no longer maintain 95% of the target force during three successive contractions. The force target and time-to-exhaustion were recorded for further analysis.

### Data analysis

All raw force signals in Newtons were filtered using a low-pass null-phase Butterworth 3 *−* order filter at a cutoff frequency of 20 Hz. The start and end of the contractions were determined when the force exceeded 10% of the maximum force measured during MVC at the beginning of the protocol. The average force of each contraction was calculated for the subsequent analysis.

#### All-out test

The maximum force *F*_max_ achieved during each contraction over the all-out test was recorded. As originally proposed (Burnley, 2009), the end-test force *EF* (conceptually related to the critical force) was defined as the average force of the last six contractions (i.e. 30 s) in the all-out test. Finally, the impulse above *EF* (*J*_EF_ = ∫ (*F*_max_(*T*) *− EF*)d*T* in N · s) was computed (Fig. 3A.).

**Figure 3.**
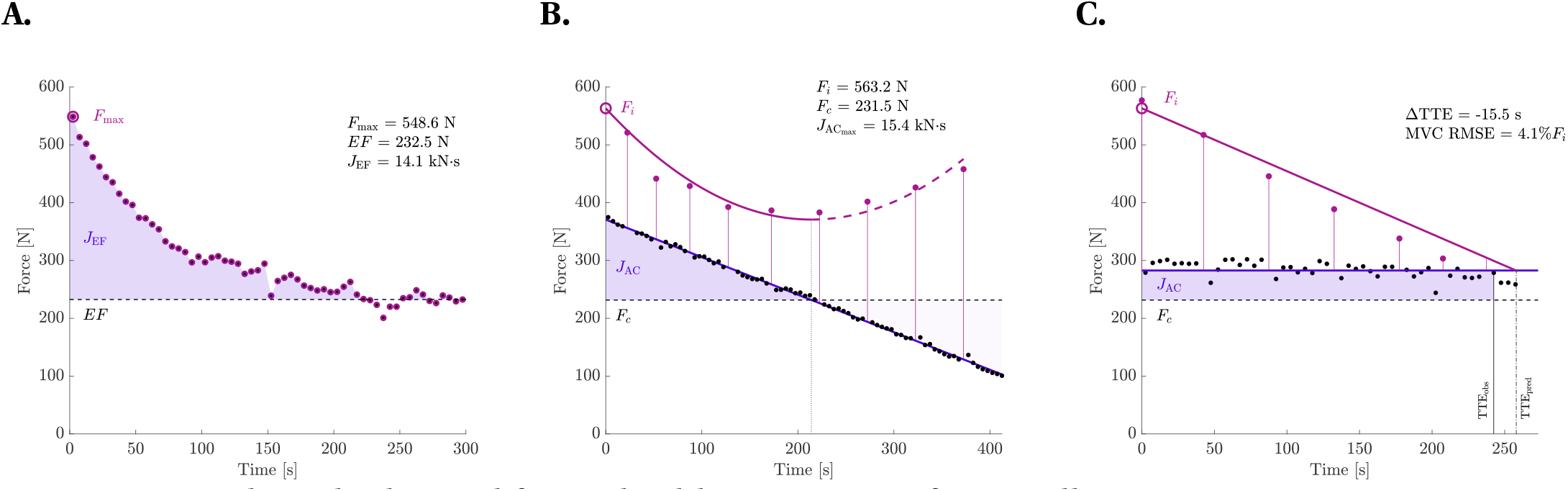
Typical result obtained for an healthy participant for (A) All-out test (B) RACLET (C) Time-to-exhaustion test. For each pannel, the maximum force obtained during a maximal voluntary contraction is represented in purple circle; the target force is represented in black dot and modelled by the blue line; the impulse above the critical force (dotted line, *EF* for the all-out test, *F*_*c*_ for the RACLET) is represented by the shaded area. The purple line represents the predicted evolution of the maximum force, initially *F*_*i*_, by the proposed model (Eq. 4). The fitted parameters for the all-out test and the RACLET are displayed on the corresponding panel (including the impulse reserve *J*_EF_ and 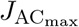 for the all-out and RACLET test, respectively). For the time-to-exhaustion test (TTE), the value of the prediction and the observation as well as the model errors are displayed (Δ TTE, MVC RMSE).

#### RACLET

To determine the values of *F*\* and *S*, all contractions except the MVCs were used to fit Eq. 3 using the least-squares minimisation method. This helped to adjust for (small) variations between the force target and what was actually performed by the participant. Next, Eq. 4 was fitted using force-time data from the MVCs performed during RACLET. Because the model used was not intended to describe the recovery phase, only data points up to three after the lowest MVC were retained. This seemed to be an acceptable trade-off between the model’s domain of application and an effective experimental means that facilitates the identification of *F*_*max*_(*t*) apex, i.e. when fatigue stops, allowing recovery. Thus, from the fitting procedure, *F*_*i*_, *F*_*c*_, and *τ* were estimated. Using these three parameters, 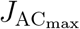 was computed as follows:

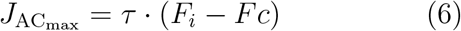

Graphically, the critical force *F*_*c*_ corresponds to the value of ramp *F* (*t*) when the minimum theoretical *F*_max_ is reached. (Fig. 3B.).

#### Time-to-exhaustion tests

Using the parameters *F*_*i*_, *F*_*c*_ and *τ* determined during the RACLET, the time-to-exhaustion for a given force target *F* was predicted for each test using Eq. 7.

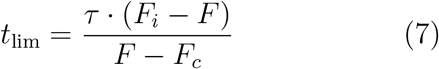

In addition to the singular moment of exhaustion, the maximum force during the test can be predicted from the generic Eq. 2 applied to the constant-force exercise. Thus, for each instant at which the maximum force was experimentally tested (i.e. every 45 s), the predicted maximum force was determined using Eq. 8.

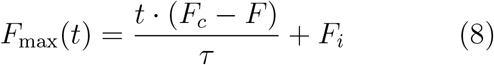

#### Statistical analysis

The data were analysed using custom-written code in MATLAB Software (2024a) and are presented as mean *±* s.d. The goodness of fit of the model from the RACLET data for each participant and session was evaluated using r^2^ as the root mean square error. In the first experiment (healthy population), the level of concurrent agreement with the proposed computational method was tested. We compared the maximal force (*F*_*i*_ vs. *F*_*max*_), critical force (*F*_*c*_ vs. *EF*), and impulse reserve (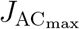 vs. *J*_EF_) parameters obtained from

RACLET versus the all-out test. The agreement between the two methods was assessed using the mean difference (i.e. systematic error; SE) and standard deviation of the differences (i.e. random error; RE) associated with their 95% confidence limits. The coefficients of correlation, r, and p *−* values were used to calculate the relationships between the two tests for each parameter. The level of statistical significance was set at p*<*0.05.

The relative and absolute reliabilities of the two RACLET were assessed only in the second experiment (i.e. the patient population). The Intraclass Correlation Coefficient (ICC2,1) (Koo and Li, 2016), Typical Error of Measurement (TEM), and Change in the Mean (CM) as well as their respective 95% confidence limits were computed for *F*_*i*_, *F*_*c*_ and 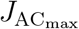. ICC and TEM were considered acceptable if they were greater than 0.7 and less than 10%, respectively, (Atkinson and Nevill, 1998). The correlation coefficient r and p *−* values were also used to calculate the correlation between the two tests.

The predictive capacity of the proposed model was tested by comparing the predicted and observed values for both time-to-exhaustion and maximal force capacity in a healthy population. Systematic and random errors, as well as the coefficient of determination r^2^, root mean square error (RMSE), and comparison between the regression equation and identity line, were computed (*vide supra*).

## Results

### RACLET feasability

For the healthy population, the RACLET goodness of fit was excellent for all participants (median r^2^ = 0.950, inter-quartile range 0.911-0.966; RMSE = 23.2 *±* 9.7 N or 3.7 *±* 1.1% when expressed relative to maximal force *F*_*i*_). The rate of perceived exertion was 14.6 *±* 2.2 (i.e. hard to very hard). A typical trace is shown in Fig. 3B.; for the patients (second experiment), the RACLET goodness of fit was very similar to that of the healthy population (median r^2^ = 0.941, inter-quartile range 0.806-0.964; RMSE = 11.8 *±* 5.5 N or 4.1 *±* 1.8% when expressed relative to maximal force *F*_*i*_).

### RACLET reliability

The intersession reliability results of the RACLET parameters for the patient population are presented in table 1. No significant differences were observed between the Test and Retest for all parameters. *F*_*i*_ and *F*_*c*_ presented a low typical error of measurement (9.2 and 8.6%, respectively), whereas it was higher for 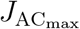 (24.0%). All ICC were *>* 0.90.

**Table 1.** Test-retest of the RACLET parameters and their reliability statistics in a population with chronic respiratory diseases.

|  | Test | Retest | CM | TEM | TEM% | ICC | r | p |
| --- | --- | --- | --- | --- | --- | --- | --- | --- |
| $F_i$ [N] | 302.93±166.20 | 284.01±148.65 | -18.92 [-34.54;-13.01] | 28.00 [19.26;51.11] | 9.2 | 0.965 [0.866;0.991] | 0.975 | <0.001 |
| $F_c$ [N] | 137.68±47.58 | 132.79±43.36 | -4.88 [-8.92;-3.36] | 11.85 [8.15;21.64] | 8.6 | 0.933 [0.768;0.983] | 0.936 | <0.001 |
| $J_{AC_{max}}$ [kN·s] | 6.48±5.01 | 6.40±5.92 | -0.08 [-1.48;-0.56] | 1.56 [1.07;2.84] | 24.0 | 0.927 [0.732;0.981] | 0.930 | <0.001 |

$F_i$ , $F_c$ and $J_{AC_{max}}$ are the initial, critical force and impulse reserve. CM: Change in the Mean; TEM: Typical Error of Measurement in absolute unit or %; ICC: Intraclass Correlation Coefficient; r and p: Pearson's correlation coefficient between Test and Retest and the associated p-value.

### RACLET validity

The results of the analysis of concurrent validity are shown in table 2, and the correlations between the methods for each parameter are displayed in Fig. 4. When comparing the all-out test and RACLET method, no statistical difference was observed for the maximal force, critical force, and impulse reserve. All RACLET parameters presented high agreement with the reference method, as shown by the lack of systematic error, all r *>* 0.90, and a random error ranging from 7 to 18%.

**Table 2.**
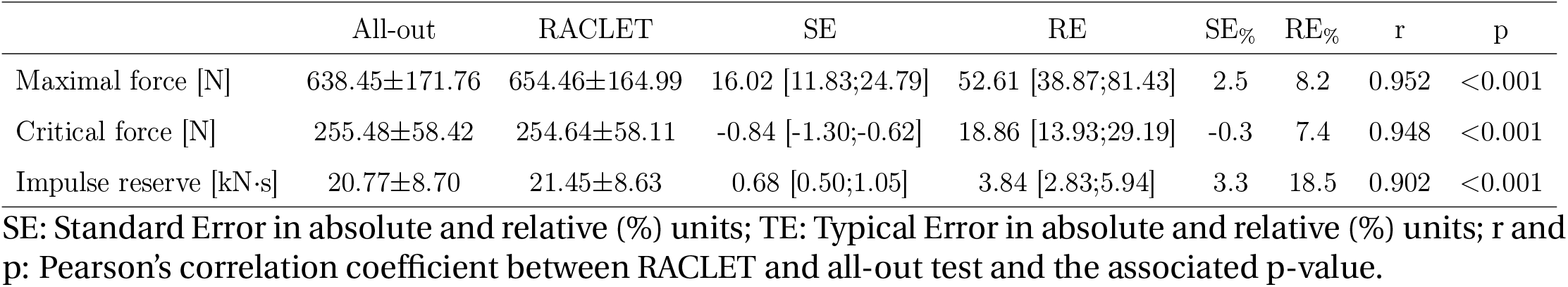
Validity statistics of the Maximal force (*F*_*i*_, *F*_max_), Critical force (*F*_*c*_, *EF*) and Impulse reserve (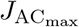, *J*_EF_) parameters determined from the RACLET compared to the all-out test.

|  | All-out | RACLET | SE | RE | SE% | RE% | r | p |
| --- | --- | --- | --- | --- | --- | --- | --- | --- |
| Maximal force [N] | 638.45±171.76 | 654.46±164.99 | 16.02 [11.83;24.79] | 52.61 [38.87;81.43] | 2.5 | 8.2 | 0.952 | <0.001 |
| Critical force [N] | 255.48±58.42 | 254.64±58.11 | -0.84 [-1.30;-0.62] | 18.86 [13.93;29.19] | -0.3 | 7.4 | 0.948 | <0.001 |
| Impulse reserve [kN·s] | 20.77±8.70 | 21.45±8.63 | 0.68 [0.50;1.05] | 3.84 [2.83;5.94] | 3.3 | 18.5 | 0.902 | <0.001 |

SE: Standard Error in absolute and relative (%) units; TE: Typical Error in absolute and relative (%) units; r and p: Pearson's correlation coefficient between RACLET and all-out test and the associated p-value.

**Figure 4.**
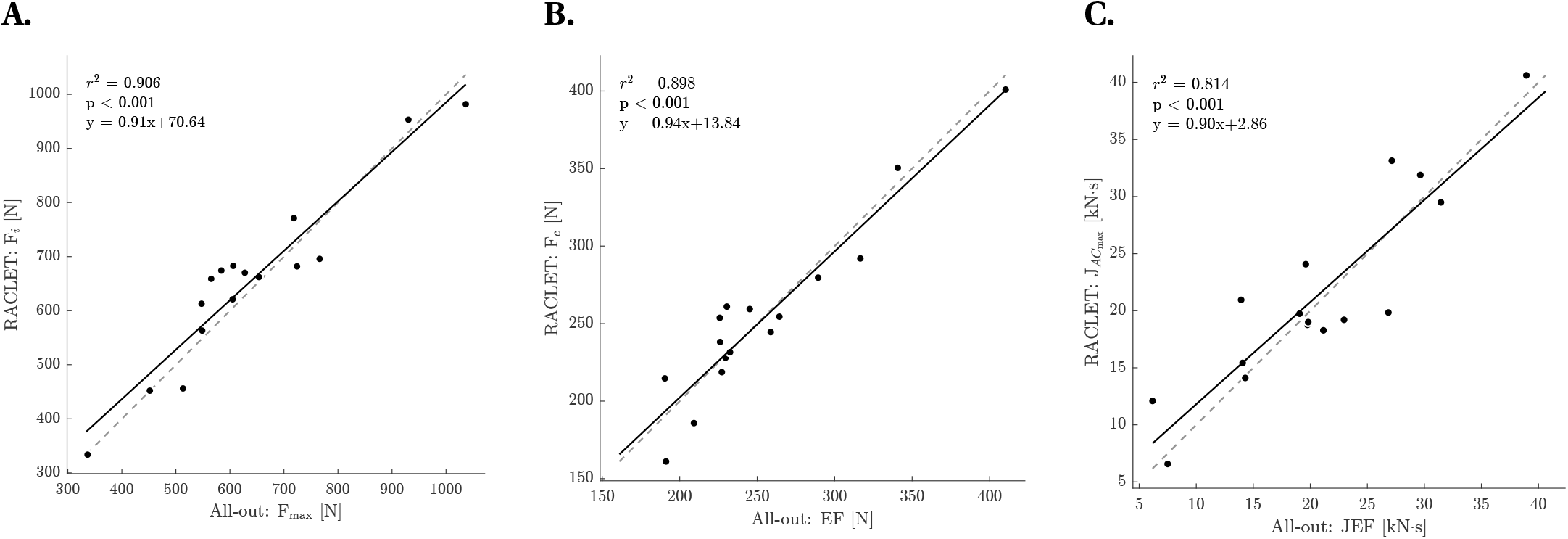
Correlation analysis of the model’s parameter obtained from RACLET and All-out method in the healthy population; (A) Maximal force; (B) Critical force; (C) Impulse reserve. Each dot represents a participant. Identity (gray dashed) and regression (black) line as well as correlation statistics and regression equation are displayed.

### Prediction capacity

The Pearson correlation analysis of the model’s predictive capacity for time-to-exhaustion (test duration) and fatigue (MVC decrease) during the test is presented in Fig. 5. For the time-to-exhaustion, r^2^ was 0.910, SE was 3.1 s (CI95% [2.56;3.88]) and RE was 52.8 s (CI95% [43.9;66.4]) for time-to-exhaustion tests of 303 *±* 171 s (RE%: 17.6%). For fatigue prediction (i.e. MVC force decrease during the time-to-exhaustion test), when pooling all tests and participants r^2^ was 0.907, SE was −1.85%*F*_*i*_ (CI95% [−2.00;−1.73]) and RE was 5.26%*F*_*i*_ (CI95% [4.91;5.67]) of maximal force. When the prediction was performed independently for each test of each participant, the prediction continued to be excellent, with a median r^2^ of 0.944 (interquartile range: 0.901-0.972) and an RMSE of 4.7 *±* 3.1% of the maximal force. A typical prediction *vs*. observed trace along a time-to-exhaustion test is presented in Fig. 3C.

**Figure 5.**
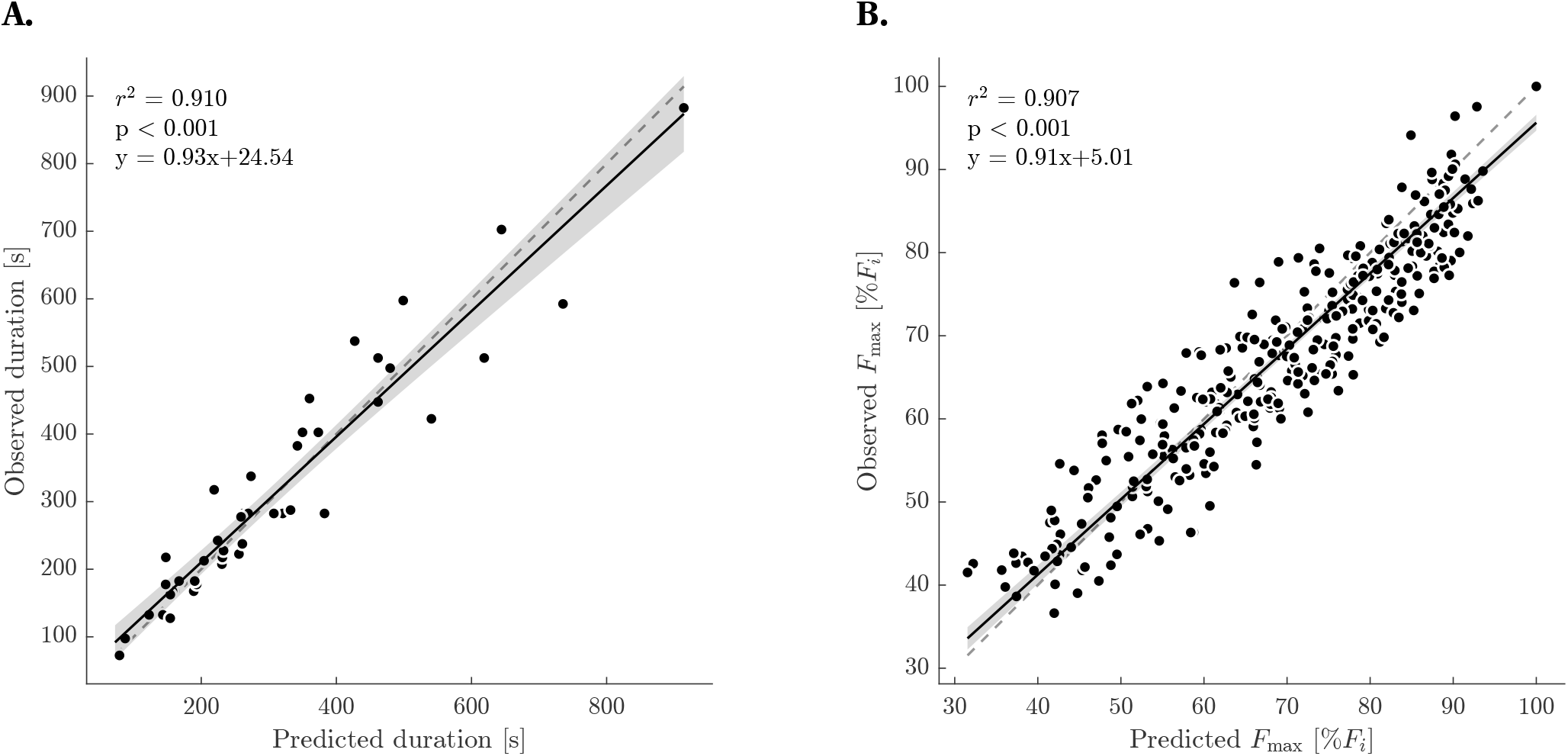
Model’s predictive capacity for (A) time-to-exhaustion (test duration) and (B) fatiguability (MVC decrease) during the test. Each dot represents either a participant (A) or a maximal voluntary contraction (B). Identity line (dashed), regresion equation, regression line (solid) and its associated 95% confidence interval (shaded area) are displayed.

## Discussion

This study aimed to propose and validate a new test, the Ramp Above Critical Level Endurance Test (RACLET), to assess the parameters of the critical force model during an isometric task without strenuous efforts. The main results were as follows: (i) the test proved easy to conduct in both healthy and pathological populations and was associated with a perception of hard but submaximal effort (RPE = 14.6 *±* 2.2); (ii) the model’s parameters demonstrated excellent reliability in patients (all ICC > 0.90); (iii) excellent concurrent validity of the critical force estimate compared to the all-out method (SE = −0.3%; RE = 7.4%); and (iv) an acceptable prediction capacity of the model for both predicting the time-to-exhaustion (RE = 56.8 s; r^2^=0.910) and the level of fatigue during a test in the severe domain (RE = 5.3%; r^2^=0.907).

First, the parameters of the proposed RACLET (total duration = 425s, decreasing ramp from 60% to 15% of the initial maximum force, MVC every 25–50s) proved effective in providing an excellent fit of the mathematical model (median *r*^2^ *≈* 0.95) while remaining submaximal. The highest RPE in the test was *≈* 15. Although this corresponds to a hard-to-very hard effort, it remains well below the RPE reported during time-to-exhaustion tests, which typically increases progressively until it reaches a value > 19 (very hard to maximal) (Nicolò et al., 2019). As the end of the test was performed below the critical force, in a state of recovery, the end-test RPE fell to *≈* 12 (light exertion). Finally, it is interesting to note that no adverse events were reported in patients with respiratory pathology, opening the door to its use in the clinical evaluation context.

Furthermore, in the clinical population tested, although the two tests were performed before and after 7 weeks of functional rehabilitation, including muscle strengthening, no significant difference was found for the three parameters, while demonstrating excellent reliability (table 1, all ICC > 0.92). This is in line with the previously reported ICC (Giles et al., 2019; McClean et al., 2023) for *F*_*c*_ (range: 0.90 - 0.95) and even better for impulse reserve (ICC range: 0.76 to 0.82). Although this was not the objective of this study, the proposed intervention did not demonstrate any benefit in terms of the force production capacity of the knee extensors. However, it is interesting to note that despite a two-month interval between the two measurements, the RACLET made it possible to characterise muscle function in terms of maximum performance (*F*_*i*_), endurance (*F*_*c*_), and exercise capacity in the severe domain 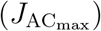 with excellent reliability.

When the parameters obtained from the RACLET were compared with those of the widely used and validated all-out method, excellent correlation (r *≈* 0.95), very low systematic error (<2.5%), and low random error (<8%) were observed for both the initial and critical force (table 2). This typical error is close to the 9% reported for the critical force of the knee extensors evaluated by the all-out method compared to the time-to-exhaustion method (Burnley, 2009). Note that some observed random errors can be attributed to the reliability of each method. Indeed, the end-test force during the all-out test typically shows a 5-10% random error (Kellawan and Tschakovsky, 2014; Starling-Smith et al., 2025). 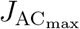 was highly correlated with *J*_EF_ but was nevertheless associated with a higher typical error (*r* = 0.90, RE = 18.5%). This is also in line with the 15% typical error and a correlation coefficient of 0.80 previously reported for the all-out vs. time-to-exhaustion method (Burnley, 2009). Indeed, the impulse reserve has been shown to be assessed with a lower signal-to-noise ratio. Therefore, the RACLET appears to provide a very good estimate of the critical force model’s parameters *F*_*i*_ and *F*_*c*_ and is acceptable for 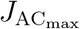 when compared to the traditional all-out method.

In addition to ensuring that parameter estimation is reliable and consistent with the reference methods, it is crucial to confirm the model’s predictive capabilities. For the critical force model, the primary assessment typically involves determining how long a constant-intensity exercise can be sustained. For each participant, we predicted the exercise duration for three different force targets higher than *F*_*c*_ with very high accuracy. Observed vs. predicted exercise time-to-exhaustion showed a very high correlation (r^2^ = 0.901) and an error of 2.5 *±* 15.6% (TE = 52.8 s for exercise duration of 303 *±* 171 s). These predictions are among the best reported in the literature, with r^2^ values usually ranging from 0.44 to 0.94 with errors from 12 to 31% (Abdalla et al., 2021, 2018; Kellawan and Tschakovsky, 2014; Vonderscher et al., 2024). Therefore, RACLET identifies a set of parameters *F*_*i*_, *F*_*c*_, and 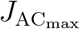 that allow the model to be applied with excellent time-to-exhaustion predictive capability, at least as good as historical methods, which are the gold standards.

Finally, an important additional value of the proposed model (Eq. 2), compared to the traditional critical power model, is its theoretical ability to describe not only the time of exhaustion (inability to sustain effort), but also fatigability (the maximal force decreases over time). A previous study revealed that once the parameters *F*_*i*_, *F*_*c*_ and *τ* were individually determined, the model proposed by Bowen et al. (2024) could accurately predict the decrease in maximum force (error: −1.8 *±* 7.8%*F*_*i*_) for a variety of exercises performed in the severe domain (i.e. above the critical intensity) during electro-induced thumb adduction (Vonderscher et al., 2024). The results of the present study (error: 1.85 *±* 5.26%*F*_*i*_; Fig. 5B.) reproduce these pilot findings and extend these observations to a voluntary task involving fatigue that may have central and/or peripheral origins (Burnley et al., 2012; Morel et al., 2019; Zarzissi et al., 2020), although this was not investigated in this study. It is interesting to note that the RACLET is precisely based on this property of the model (the description of the evolution of maximum force during exercise) to determine the parameters without requiring the strenuous efforts required in multiple time-to-exhaustion or all-out methods.

The performance of this test, coupled with its feasibility (limited fatigue, simple instructions, short duration), opens up numerous practical applications hitherto impossible with traditional time-to-exhaustion and all-out methods. Indeed, in the context of whole-body endurance training, whether for patients or athletes, individualisation based on a physiological threshold (e.g. critical power, gas exchange threshold) yields better results than relying on an arbitrary percentage of maximal capacity (Meyler et al., 2025; Vallet et al., 1997). For this reason, learned societies have focused on COPD advocates for reconditioning based on the metabolic threshold (Maltais et al., 2014). However, the latter approach is not available for localised muscle training, which offers yet another benefit of reducing strain on the cardiorespiratory system and thereby aids in preventing dynamic hyperinflation in this population (Broxterman et al., 2020; O’donnell et al., 2001). Therefore, the RACLET could be used to propose individualised muscle physiological threshold values for tailored training sessions in a rehabilitation context. However, the benefits of RACLET cannot be confined to assessing and individualising physical activity programs for vulnerable populations. All conditions requiring a large number of critical force measurements (e.g. longitudinal follow-up, address research questions imposing a very large number of experimental conditions), or simply to reduce the cost of participating in a study for ethical reasons, will benefit from the RACLET.

Nevertheless, it is crucial to bear in mind that the very good reliability and validity results presented in this study are directly linked to the quality of the target force reaching, the maximum voluntary force measurements, and the set of RACLET parameters (*F*\*, slope *S*, and test duration). Anyone wishing to implement this test should pay particular attention to these factors. Typically, an incorrectly configured test (too light or too hard) makes it impossible to exploit the results or complete the test. To help anyone wishing to set up the RACLET, all the procedures for designing and analysing the test are described and freely available in Excel and Matlab versions on the FoVE research group’s Gitlab (https://gricad-gitlab.univ-grenoble-alpes.fr/fove/methods/raclet).

## Conclusion

Based on a mathematical model previously developed by our team, this study proposes and validates a new test: the Ramp Above Critical Level Endurance Test. The parameters of the model are the initial maximum force *F*_*i*_, critical force *F*_*c*_ and characteristic time *τ* (from which we can also determine 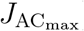), which proved to be reliable, valid, and predictive as gold standard methods. This opens up new opportunities for critical force assessment, both in diagnosis and training guidance for patients and athletes, as well as in research, to remove the major bottleneck that is the difficulty and/or time-consuming nature of gold standard methods.

## Abbreviations

COPD: Chronic obstructive pulmonary disease
CM: Change in the Mean
*F*_*c*_: Critical theoretical force
*F*_*i*_: Initial maximal theoretical force
*F*_max_: Maximal theoretical force
*F*^*⋆*^: Force at the start of RACLET ICC Intraclass Correlation Coeficient
*J*_AC_: Impulse done Above Critical Force
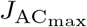 Maximal *J*_AC_: that can be done *J*_EF_ Impulse done Above End force
RACLET: Ramp Above Critical Level Endurance Test
RE: Random Error
RMSE: Root mean square error
*S*: Force ramp slope of RACLET
SE: Standard Error
*τ*: Typical time of fatigability
*t*_*c*_: time at which critical power is reached
TEM: Typical Error of Measurement
*t*_lim_: Duration of Time to exhaustion

## Acknowledgments

Support was provided by the French National Agency [ANR-22-CE14-0073] to B. Morel. The authors declare no conflict of interest relevant to this article. The results of this study are presented clearly and honestly, and without fabrication, falsification, or inappropriate data manipulation. We would like to thank P. Rozier-Delgado for his careful proofreading of the manuscript.

